# Seasonal cycles in a seaweed holobiont: A multiyear time series reveals repetitive microbial shifts and core taxa

**DOI:** 10.1101/2024.10.23.619769

**Authors:** Chantal Marie Mudlaff, Florian Weinberger, Luisa Düsedau, Marjan Ghotbi, Sven Künzel, Guido Bonthond

## Abstract

Seasonality is an important natural feature that drives cyclic environmental changes. Seaweed holobionts, inhabiting shallow waters such as rocky shores and mud flats, are subject to seasonal changes in particular, but little is known on the influence of seasonality on their microbial communities.

In this study, we conducted a bi-monthly, three-year time series to assess the seasonality of microbial epibiota in the seaweed holobiont *Gracilaria vermiculophylla*. Our results reveal pronounced seasonal shifts that are both taxonomic and functional, oscillating between late winter and early summer across consecutive years. While epibiota varied taxonomically between populations, they were functionally similar, indicating that seasonal variability drives functional changes, while spatial variability is more redundant.

We also identified seasonal core microbiota that consistently (re)associated with the host at specific times, alongside a permanent core that is present year-round, independent of season or geography. These findings highlight the dynamic yet resilient nature of seaweed holobionts and demonstrate that their epibiota undergo predictable changes. Therewith, the research offers important insights into the temporal dynamics of seaweed-associated microbiota, and demonstrates that the relationship between seaweed host and its epibiota is not static, but naturally subject to an ongoing seasonal succession process.

## INTRODUCTION

Seasonality is a global environmental feature which plays an important role in structuring communities in terrestrial and aquatic environments (White & Hastings 2020). To communities, seasonality is a natural source of variability, characterized by cyclic changes in temperature and photoperiod, but also by other variables such as salinity, rainfall, phytoplankton blooms, anthropogenic stressors, upwelling, and nutrient pulses (Lisovski *et al*. 2017). Seasonal fluctuations are also prevalent in free-living microbial communities (Fuhrman *et al*. 2015; Gilbert *et al*. 2012) and microbial communities associated with various hosts (Sharp *et al*. 2017; Ferguson *et al*. 2018; Gobbi *et al*. 2020; Risely *et al*. 2021), including seaweeds (Bengtsson *et al*. 2010; Park *et al*. 2022; Tujula *et al*. 2010).

Seaweeds are typically found in coastal habitats, such as rocky shores and intertidal mudflats, which experience strong seasonal forces (Benincà *et al*. 2015). Naturally, seaweed holobionts are surrounded by microbial life. The algal surface is in direct contact with the surrounding water and acts as substrate on which microorganisms can settle (Wahl *et al*. 2012). These colonizing communities are typically dominated by bacteria, but also include microalgae, fungi, protists, and viruses (Egan *et al*. 2013; Van Der Loos *et al*. 2019). On the interface between the seawater and the inner tissue of the seaweed, epibiota form a dynamic biofilm, which has also been termed the “second skin” and influence the host physiologically, chemically and biologically (Wahl *et al*. 2012). These epibiota include (opportunistic) pathogens, but also beneficial microbes that promote the host’s development and fitness, such as e.g., sporulation (Weinberger *et al*. 2007) or morphogenesis (Weiss *et al*. 2017, reviewed in Egan *et al*. 2013), pathogen recognition, and chemically mediated defense mechanisms (Li *et al*. 2022; Longford *et al*. 2019; Rao *et al*. 2007; Saha & Weinberger 2019).

Epiphytic communities are complex and their structure depends on host morphology (Lemay *et al*. 2021), varies by species (Lachnit *et al*. 2009, 2011) or even lifecycle stage (Lemay *et al*. 2018; Bonthond *et al*. 2022), and differs along the algal thallus (Paix *et al*. 2020), implicating the specific and diverse niches provided by the host and the microbial partners. The holobiont is exposed to variable environmental conditions, which may alter the epibiota composition either directly, or indirectly via host physiological responses to the changing environment. For instance, seaweed epibiota have been found to strongly vary with salinity (Stratil *et al*. 2014; Van Der Loos *et al*. 2023) and temperature (Stratil *et al*. 2013; Bonthond *et al*. 2023). Thus, seaweed epibiota are shaped by a combination of host and environment. As both environmental conditions and host physiology are seasonal, it is not surprising that seasonal patterns have been detected in seaweed epibiota (Bengtsson *et al*. 2010; Park *et al*. 2022; Tujula *et al*. 2010). While it is informative to document microbial changes within the holobiont from one season to another, it is important to distinguish which changes are repetitive. Such cyclic patterns reflect an ongoing interaction between host and microbiota, with long multigenerational histories. Microbial taxa that are present irrespective of season, or that return interannually, may represent important core symbionts (beneficial or harmful). Studying holobionts during different seasons across several years may resolve such permanent and/or seasonal core microbiota and therewith contribute to the identification of important host-microbe associations and to better understand the complex dynamics within the holobiont.

*Gracilaria vermiculophylla* is a well-studied holobiont. This perennial rhodophyte is native to the North-West Pacific (Kim *et al*. 2010; Krueger-Hadfield *et al*. 2017, 2021) but has become invasive across the Northern Hemisphere, including the Eastern Pacific southward to Mexico (Bellorin *et al*. 2004), the North American coasts in the Western Atlantic (Freshwater *et al*. 2006; Thomsen *et al*. 2006), as well as the European coasts at the Eastern Atlantic, extending towards northern parts of the North Sea and the South Western Baltic Sea (Rueness 2005; Thomsen 2007; Weinberger *et al*. 2008). Epibiota associated with *G. vermiculophylla* have been studied across the Northern Hemisphere (Bonthond *et al*. 2020), which revealed that some epi- and endobiota were part of a core, i.e., a group of microbial taxa that was associated with the *G. vermiculophylla* holobiont irrespective of the host geography. This holobiont was also studied in controlled experiments in the lab, and its microbiota were sampled repeatedly over several weeks to months (Bonthond *et al*. 2021a, 2023). These time series demonstrated that epi- and endobiota within the holobiont have strong temporal variation and that not all geographic core microbiota were temporally stable. Altogether, this may indicate that many core microbes are rather season specific. Given the wide geographic stretch, across which spatial variability and core microbes have already been studied in Bonthond *et al*. (2020), *G. vermiculophylla* presents a suitable seaweed holobiont to characterize seasonal variability and core microbiota.

The aim of the present study was therefore to evaluate seasonal variability in *G. vermiculophylla* associated epibiota as well as to characterize cyclic patterns and permanent core microbiota (i.e., seasonal and interannual). To this end, we conducted a bi-monthly time series sampling of *G. vermiculophylla* individuals from two distinct populations over three consecutive years, resulting in a dataset with 18 repeated measures. We hypothesized that prokaryotic epibiota associated with *G. vermiculophylla* show seasonality, in terms of taxonomic and functional composition and in terms of diversity. Furthermore, we hypothesized that *G. vermiculophylla* harbors core microbiota, which are (i) permanent (i.e., associated irrespective of time and space) as well as (ii) season–specific (i.e., consistently present in the holobiont during specific times of the year).

## EXPERIMENTAL PROCEDURES

### Sample collection

Seaweeds were collected in Nordstrand (Germany) at the North Sea (54°29’9.34”N 8°48’44.65”E) and in Heiligenhafen (Germany) at the Baltic Sea (54°22’46.7”N 10°58’57.5”E; Fig. 1A-D). These populations were chosen based on their distinct environmental features. The North Sea population is found in the intertidal zone. Here, the perennial *Gracilaria vermiculophylla* occurs mainly attached to hard substratum and can build massive mats during spring and/or summer. In contrast, the Baltic Sea population is situated in a small lagoon sheltered from turbulences and is only experiencing wind driven sea level fluctuations. Here, the *G. vermiculophylla* individuals are not attached to substratum but rather loosely embedded in the soft sediment. During winter, the number of individuals typically reduces substantially and sometimes appears to be absent. Whereas individuals in the North Sea population are exposed to fully marine salinities (i.e., ∼ 24 to 32, Fig. 1E) and diurnal air exposure, the Baltic Sea population experiences rather brackish salinities (between ∼ 10 and 20, Fig. 1E) as well as less and irregular air exposure. Generally, the pH is more similar between the two populations, normally fluctuating between 7 and 9, although an outlier was detected in Heiligenhafen of 5.3, recorded during early summer in year 1 (Fig. 1F).

**Figure 1.**
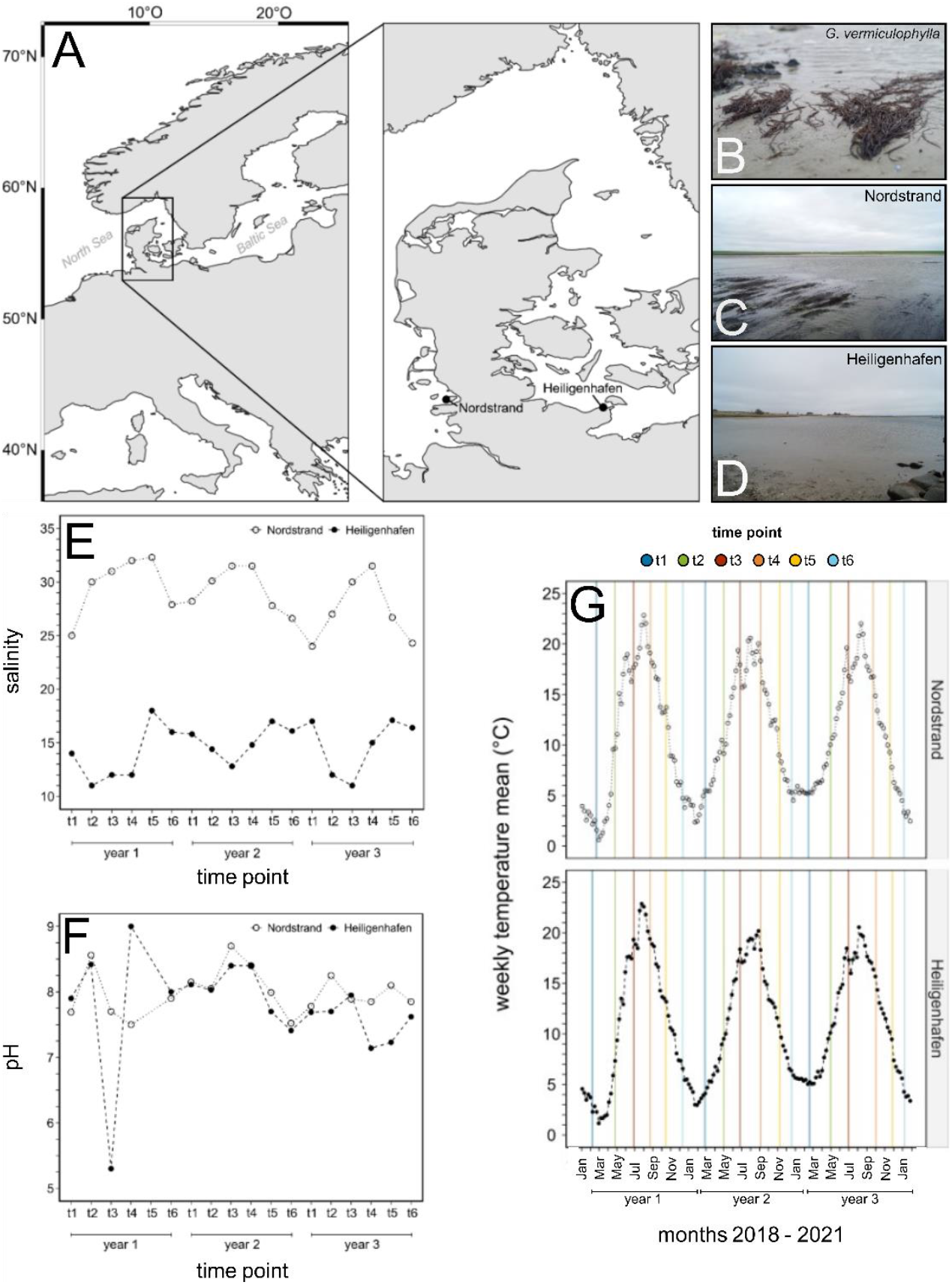
Overview of the two collection sites Nordstrand (North Sea) and Heiligenhafen (Baltic Sea), where the bi-monthly sampling of *Gracilaria vermiculophylla* was carried out, and environmental parameters during the three-year time period. (**A**) Map showing the two collection sites. (**B**) Habitus of *Gracilaria vermiculophylla*. (**C**) North Sea population found at Nordstrand, (**D**) Baltic Sea population found at Heiligenhafen. (**E**) Salinity and (**F**) pH measured during the field sampling. (**G**) The weekly water temperature (°C) obtained from nearby measuring stations, with vertical lines depicting the exact collection time points. Temperature data provided by the German Federal Maritime and Hydrographic Agency (BSH, 2023).

The sample collection took place in bi-monthly intervals covering a three-year time period from February 2018 to January 2021. During these years water temperatures at locations nearby to Nordstrand and Heiligenhafen oscillated similarly between minima of 1 to 6°C in winter and maxima of 21 to 23°C in summer (Fig. 1G). At each sampling point, 10 *G. vermiculophylla* individuals were collected with gloves and placed separately into plastic bags. To avoid collection of the same individual, the individuals were sampled at least 1 m away from each other. In the North Sea population, only attached algae were sampled. Additionally, three 50 ml water and two 15 ml sediment samples were taken at both sites. Subsequently, around 0.25 ml sediment was transferred into a 2 ml tube containing absolute ethanol. After collection, all samples were transported in a cooling box back to the facilities of GEOMAR Helmholtz Centre for Ocean Research in Kiel (Germany) where they were stored at 4°C and processed within two days maximum. In the laboratory, salinity and pH were measured for both collection sites from one of the three water samples. The remaining water samples were processed further together with the seaweed samples.

### Generating epiphytic extracts

To generate extracts from the prokaryotic epibiota associated with *G. vermiculophylla*, the method in Bonthond *et al*. (2020) was followed. In brief, a branch of 1 ± 0.25 g was transferred into a 50 ml tube. Approximately 10 glass beads and 15 ml artificial seawater of the respective salinity (prepared from distilled water and sodium chloride) were added. Besides the field samples, at least one blank was prepared for each sampling event, containing only glass beads and distilled water. Afterwards, all samples were vortexed for 2 min at maximum rotation speed. After vortexing, the algal tissue was removed. For the samples collected in the first sampling year, the epiphytic suspension was filtered as in Bonthond *et al*. (2020). During the second and third year, the centrifugation method of Ficetola *et al*. (2008) was used. In brief, 33 ml absolute ethanol and 1.5 ml sodium acetate were added to 15 ml epiphytic suspension, water, and blank samples. All tubes were mixed and either processed immediately or cooled at 4°C and processed in the next days. The 50 ml tubes were then centrifuged for 10 min at 14,000 g. The supernatant was discarded and the pellet was preserved in 1 ml absolute ethanol. The generated epiphytic algal extracts (on filters or in 1 ml ethanol), water, sediment, and blanks were stored at – 20°C until DNA extraction.

### DNA extraction & amplicon library preparation

Ethanol was removed from the samples by evaporation in a vacuum centrifuge for at least 1 hour at 45°C. If the evaporation of the alcohol was unsuccessful after several hours, the remaining ethanol was removed by lyophilization. Filters were fragmented to small pieces with sterile scissors. Subsequently, DNA was extracted following the Cetyltrimethylammonium bromide (CTAB)–chloroform protocol from Doyle & Doyle (1987). The amplicon library was prepared following a two-step PCR approach by Gohl *et al*. (2016), using the same indexing primers and KAPA HIFI HotStart polymerase (Roche). The first PCR targeted the V4 region of the 16S rRNA gene using the forward primer U515F (S-*-Univ-0515-a-S-19) and the reverse primer 806R (S-D-Arch-0786-a-A-20; Klindworth *et al*. 2013) with adapters for the second PCR on 5’ ends. The first PCR program began with a step of 5 min at 95°C and was followed by 25 cycles of denaturation for 20 sec at 98°C, annealing for 15 sec at 55°C, and elongation for 1 min at 72°C. For water samples which were not successfully amplified, 30 cycles were used in a repeated PCR attempt.

For the second PCR, the amplicon products were diluted 1:10 and used as template. PCR was conducted following the same cycling program, but with 10 cycles of denaturation and an additional final elongation step of 10 min at 72°C. Subsequently, PCR products were visualized by gel electrophoresis and relative amplicon concentrations were estimated from gel pictures using the software Image J Fiji (Fiji for Mac OS X Version 1.0) to accordingly adjust volumes in the library pooling. The pooled library was purified with a gel extraction step by using the ZymoClean Gel DNA recovery kit (ZymoResearch) following the supplied protocol, quantified with qPCR, and sequenced as paired-end reads (2 × 300) on the Illumina MiSeq platform at the Max Planck Institute in Plön (Germany).

### Data processing

A total of 406 samples (including controls) was processed with the software Mothur (v.1.43.0 and v.1.45.3; Schloss *et al*. 2009) following an inhouse script. Accordingly, unique sequences were aligned to and classified with the SILVA reference alignment v132 (Quast *et al*. 2013). The sequences were open-reference clustered to the 3% OTUs from a field study by Bonthond *et al*. (2020) with the *cluster*.*fit()* function (Sovacool *et al*. 2022). Sequences of mitochondrial, chloroplast, eukaryotic, and unknown origin were removed. Finally, OTUs that were singletons in the full dataset and samples with < 1000 reads were removed. The raw demultiplexed amplicon reads were deposited in the SRA database (accession: PRJNA1155875). Predicted metagenomes based on KEGG Ortholog (KO) annotations (Kanehisa *et al*. 2014) were obtained with PICRUST2 v2.5.0 (Douglas *et al*. 2020).

The variability in the taxonomic (based on OTUs) and functional (based on predicted KOs) community composition over time was visualized with non-metric multidimensional plots (nMDS) using Bray-Curtis distances with the R package vegan (Oksanen 2010), based on rarefied data of either all samples including all substrates (alga, water, and sediment) or solely algal substrates only. Additionally, trajectories were drawn chronologically through the group centroids of algal samples from the same sampling event. To test for differences in taxonomic and potential functional community composition, permutational multivariate analysis of variance (PERMANOVA) was applied on the unrarefied OTU data set, by using 9,999 permutations in the R package vegan (Oksanen 2010; usage of the adonis2 function). First, a PERMANOVA was run on the full OTU and KO datasets, including alga, water, and sediment samples. The model included the variables season (as factor with 6 levels), year (as factor with 3 levels), population (as factor with 2 levels), substrate (alga, water, and sediment), and all interactions. Subsequently, a PERMANOVA was run on the OTU and KO datasets of only algal samples, with the variables season, year, population, and all possible interactions. In both PERMANOVAs, the sequencing depth was logarithmic transformed (LSD) and included as covariate to account for its effect.

To analyze the microbial diversity the asymptotic species richness (S_Chao_) was calculated with the R package iNEXT v3.0.0 (Chao *et al*. 2014; Hsieh *et al*. 2016). Second, as a measure of evenness, the probability of interspecific encounter (PIE; Hurlbert 1971), was calculated with the R package mobr v2.0.2 (McGlinn *et al*. 2019). For both diversity measures the total data set containing exclusively algal OTU read counts was used. For the S_Chao_, a generalized linear model (GLM) was fitted, including the main effects of season (as factor with 6 levels, corresponding to the sampling events repeated over three years), year (as factor with 3 levels), population (as factor with 2 levels), and all possible interactions. The model assumed a gaussian family distribution, with a logarithm in the link function. The log transformed sequencing depth (LSD) was included as a covariate to account for variation in total read counts across samples. For the PIE, a GLM with the same model structure was used. PIE was logit transformed to meet the model assumptions. As the sequencing depth was not significant, it was excluded from the model. Analysis of variance (ANOVA) was applied to the GLM of S_Chao_ and PIE to test for significance. If the main effects season, year, and population were significant, post-hoc analysis was performed by pair-wise comparisons in both models on all possible interactions between the main effects by using the R package emmeans v1.8.9 (Lenth 2022).

### Defining permanent and seasonal core microbiota

Core microbiota were identified using two alternative compositional approaches by identifying differentially abundant OTUs (see Shade & Handelsman 2012 for definitions on core microbiomes). With both approaches we defined both a permanent core, including OTUs persistently detected within the epibiota across all seasons and years, and two seasonal cores, including OTUs consistently detected within either summer or winter.

The first approach was based on multivariate GLMs (mGLMs) from the R package mvabund v4.2.1 (Wang *et al*. 2012), which was also used to characterize the spatial core in Bonthond *et al*. (2020). mGLMs were fitted on the cumulative 95% most abundant OTUs with > 25% prevalence and assumed a negative binomial distribution. For the permanent core, the variables substrate (alga, water or sediment), season (6 levels, t1:t6) and year (3 levels) were included as predictors. For seasonal mGLM cores, the OTU matrix was reduced to season time points t1 (late winter), t3 (early summer), t4 (late summer) and t6 (early winter) and the factor ‘season’ was reduced to only two levels representing the seasonal extremes (winter: t6 & t1, summer: t3 & t4), and included in the model together with the factor year. Both mGLMs included the LSD as offset to correct for different sequencing depths across samples. Models were resampled using the summary.manyglm function with 500 bootstrap iterations, which were restricted within populations. The p-values were obtained through Wald tests. OTUs were considered part of the permanent core when the coefficients substrate_alga:water_ and substrate_alga:sediment_ were negative (reflecting higher relative OTU abundances associated with algal samples compared to water and sediment) and with corresponding p-values < 0.01. Similarly, OTUs with positive and negative coefficients for the factor season_summer:winter_ and with p-values < 0.01, were considered winter and summer core OTUs, respectively.

In addition to the mGLM core, we also defined a compositional core with the linear discriminant analysis effect size method (LEfSe, Segata *et al*. 2011) through the online interface at the webpage from the Huttenhower lab (https://huttenhower.sph.harvard.edu/galaxy/; accessed May 2023). For the permanent LEfSe core, OTUs significantly more abundant in epibiota samples compared to both water and sediment samples were considered core OTUs. Also, for seasonal LEfSe cores, the dataset was reduced to summer (time points t3 and t4 combined) and winter (time points t1 and t6 combined) to identify OTUs significantly more abundant in either season.

## RESULTS

### Sequencing summary

After all quality filtration steps, the final dataset counted 262 samples (170 algal, 29 water, and 63 sediment samples) and 14,874,961 sequencing reads, clustered into 45,751 OTUs, including 17,670 OTUs that were already identified in Bonthond *et al*. (2021). The overall most abundant OTU in the holobiont was classified to *Granulosicoccus* (OTU97) with a mean relative abundance of 4.93% and 99.4% occupancy, followed by OTUs classified to Alphaproteobacteria (OTU12, 1.82% abundance and 100% occupancy), Rhodobacteraceae (OTU00003, 1.73% relative abundance and 100% occupancy), and *Desulforhopalus* (OTU1577, 1.51% relative abundance and 91.60% occupancy). The most abundant families were Rhodobacteraceae (18.17%), Flavobacteriaceae (15.13%), Saprospiraceae (10.73%) and Thiohalorhabdaceae (4.81%, Fig. 2).

**Figure 2.**
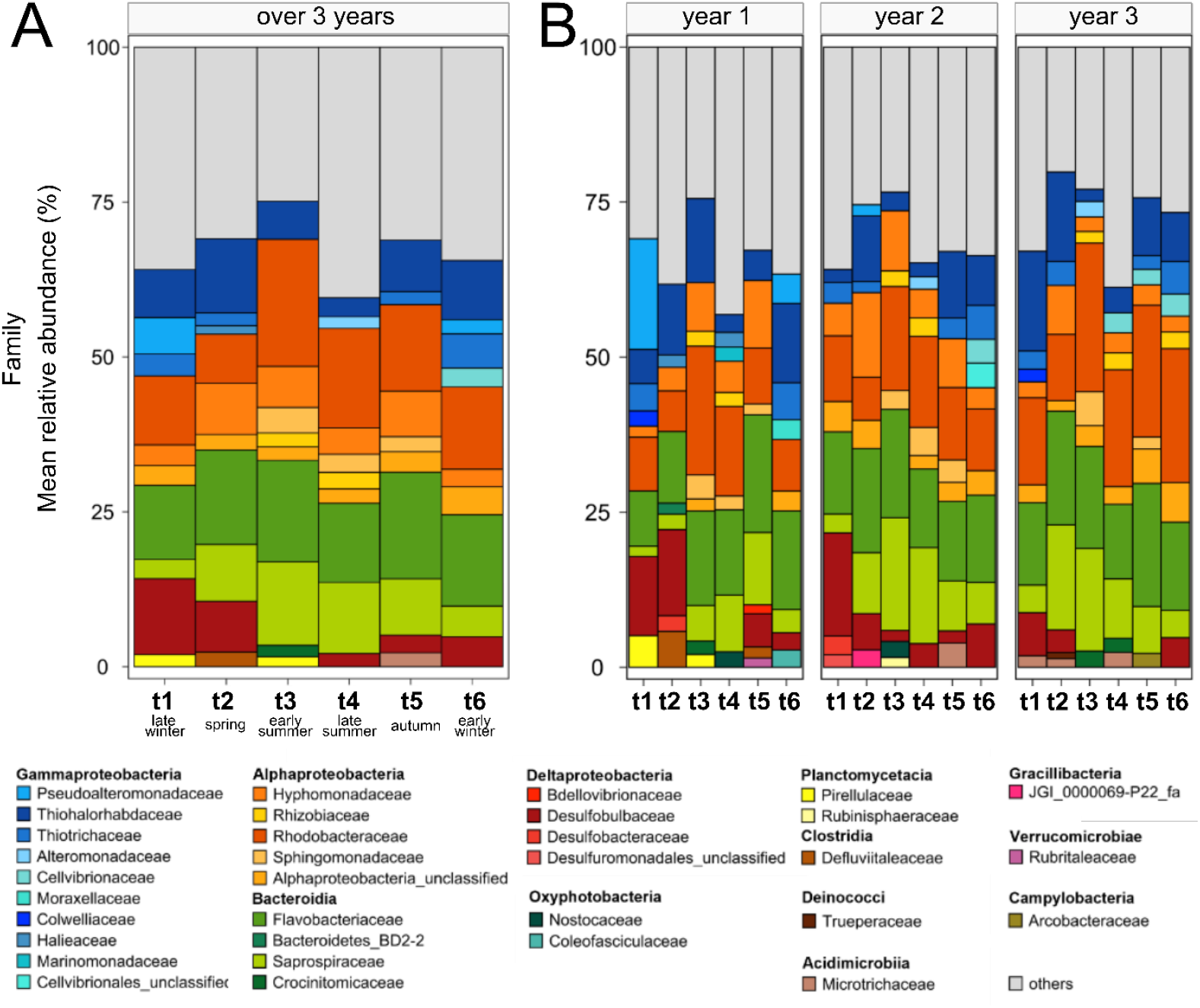
Microbial community composition of the ten most abundant families associated with the surface of the red seaweed *Gracilaria vermiculophylla*, at (**A**) season time points averaged over populations and years, and (**B**) season time points for each year averaged over populations. Shown is the mean relative abundance in percent (%).

The collected prokaryotic surface communities associated to the red seaweed *Gracilaria vermiculophylla* compositionally differed from the seawater and sediment communities. Although the classes Gammaproteobacteria, Alphaproteobacteria, Bacteroidia, and Deltaproteobacteria contributed to the general community composition structure and were shared between all three substrates, detectable differences in abundance occurred already in the top 10 families (Fig. S1). Notably, the ten most abundant prokaryotic families contributed around 65% of all families present on *G. vermiculophylla*, but less than 50% of those present in sediment and water. The families Flavobacteriaceae and Rhodobacteraceae were particularly important in algal and seawater samples, while Halieaceae, Desulfobulbaceae, Desulfobacteraceae, Chromatiaceae, and Pirellulaceae were more important in sediment samples. Interestingly, Thiohalorhabdaceae, Thiotrichaceae, Sphingomonadaceae, and Pseudoalteromonadaceae only occurred in the top 10 families of algal samples (Fig. S1).

Microbiota on algal surfaces varied considerably over time. Regarded for both populations together and throughout the years, Gammaproteobacteria were mainly represented by Thiohalorhabdaceae, Alphaproteobacteria by Rhodobacteraceae, and Bacteroidia by Flavobacteriaceae and Saprospiraceae. This pattern was best visible in the second and third year (Fig. 2). Additionally, each year had a specific pattern of interchanging families along seasons. Desulfobulbaceae tended to be less prominent in summer than in winter, while an opposite pattern was observed for Rhizobiaceae (Fig. 2). However, exceptions occurred. For instance, Rhizobiaceae were also relatively abundant in winter (t6) in the third year and an exceedingly high abundance of Pseudoalteromonadaceae was once observed at t1 in the beginning of the first year.

Microbiota on algal surfaces also varied between the populations (Fig. S2). Thiohalorhabdaceae, Thiotrichaceae, Microtrichaceae, and Rhizobiaceae solely occurred in the ten most abundant families at Nordstrand, whereas dominant families associated primarily to Heiligenhafen were Pseudoalteromonadaceae, Sphingomonadaceae, Cellvibrionaceae, and Pirellulaceae. A comparison of community compositions at the two populations over time (Fig. S3) generally confirmed these differences, as well as the pertaining dominance of a small group of families within each population. At the same time, the clear dominance of certain microbial families at specific seasons mainly disappeared and a more fine-tuned picture established. More families appeared within the top 10, which often exhibited more fluctuating abundances throughout the year. Families that emerged predominantly in Nordstrand were for instance Bdellovibrionaceae and JGI_0000069-P22_fa (Gracillibacteria). In contrast, new families belonging to the Oxyphotobacteria appeared in Heiligenhafen among the top 10 mainly in summer (t3) and beginning of autumn (t4) throughout the first two years (Fig. S3).

### Community composition

An nMDS based on taxonomic composition clearly separated algal samples and sediment samples, while seawater samples arranged amid those two (Fig. S4). A PERMANOVA confirmed that much compositional variation was explained by the substrate (R^2^ = 0.117; p < 0.001, Table S1). Among algal samples only, another significant source of differences in microbial taxonomic composition was the population (PERMANOVA, R^2^ = 0.053; p < 0.001, Table S1; for algal samples only R^2^ = 0.109; p < 0.001; Table S2). nMDS correspondingly separated samples collected from algal surfaces and sediment samples at Nordstrand and Heiligenhafen nearly completely, while water samples exhibited more compositional similarity between populations (Fig. S4).

Microbial taxonomic community composition varied significantly over time. Season and year together (individually or in interaction with other factors) explained much of the compositional differences between samples (PERMANOVA, Table S1 for all samples and Table S2 for algal samples only). nMDS indicated that the taxonomic composition of epiphytic communities varied by season, with similar seasonal shifts across years (Fig. 3A). Correspondingly, PERMANOVA confirmed that the clustering by season (R^2^ = 0.092; p < 0.001; Table S2, Fig. 3B) was stronger than clustering by year (R^2^ = 0.028; p < 0.001; Table S2). Despite substantial differences between the two populations (R^2^ = 0.109; p < 0.001) the seasonal shifts in microbial composition were similar in direction and magnitude.

**Figure 3.**
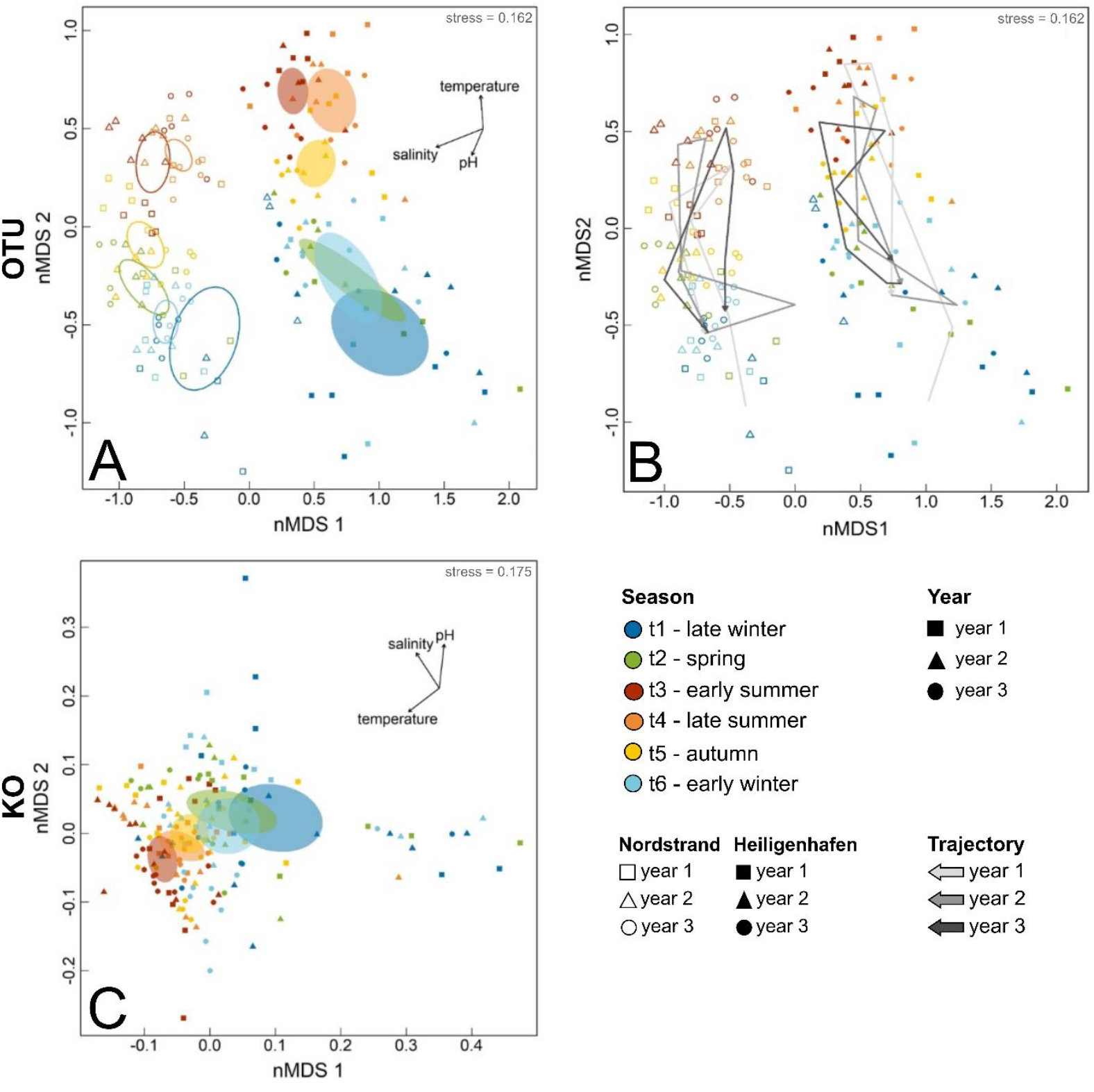
Nonmetric-multidimensional scaling (nMDS) of the microbial taxonomic (**A, B**) and functional (**C**) diversity associated with the red seaweed *Gracilaria vermiculophylla*. The season time points are pooled over populations and years and are displayed with a unique color coding as follows: t1 (late winter) in dark blue, t2 (spring) in green, t3 (early summer) in red, t4 (late summer) in orange, t5 (autumn) in yellow, and t6 (early winter) in light blue. The corresponding ellipses are represented with a 95% confidence interval. The years can be differentiated by their shape i.e., a square for year 1, triangle for year 2 and dot for year 3. Additionally, the abiotic factors temperature, salinity, and pH are plotted. The stress value is given in the upper right corner. The taxonomic (based on OTUs) and functional (based on predicted KOs) diversity shown is based on rarefied data including solely algal samples.

While functional community composition was also strongly shaped by season (R^2^ = 0.092; p < 0.001), differences between populations were limited (R^2^ = 0.007, p = 0.041) and not clearly visible in the nMDS, although functions are derived from taxonomy. Also, the factor year explained more functional diversity (R^2^ = 0.016; p = 0.008) than the factor population, which was not the case for taxonomic diversity (Tables S2 and S3). nMDS confirmed that functional diversity oscillated between summer and winter (Fig. 3C)

### Diversity

The asymptotic OTU Chao richness (S_Chao_) of microbial communities on the surface of *G. vermiculophylla* showed significant seasonal variation (Likelihood Ratio χ^2^(5) = 59.076, p < 0.001; Table S4). It was characterized by a minimum in early summer (t3) and a maximum in late winter (t1) and was rather constant from summer (t4) to early winter (t6) (Fig. 4A; Table S4). A different seasonal pattern emerged for evenness (Likelihood Ratio χ^2^(5) = 86.574, p < 0.001; Table S5). PIE was highest in summer (t4) and minimal during winter (t6 and t1, Fig. 4B). Similar to Chao richness in OTUs, Chao richness of predicted functions showed strong seasonal variation (Likelihood Ratio χ^2^(5) = 34.03, p < 0.001; Table S4) with a maximum in late winter (t1) and minimum in early summer (t3), but transitioned more gradually towards these extremes over the other seasonal time points (Fig. 4C, Table S4). Evenness in predicted functions varied with season as well (Likelihood Ratio χ^2^(5) = 39.519, p < 0.001; Table S5), but yielded a more complex trend with multiple optima (t4, t6) and minima (t2, t3, t5, Fig. 4D, Table S5).

**Figure 4.**
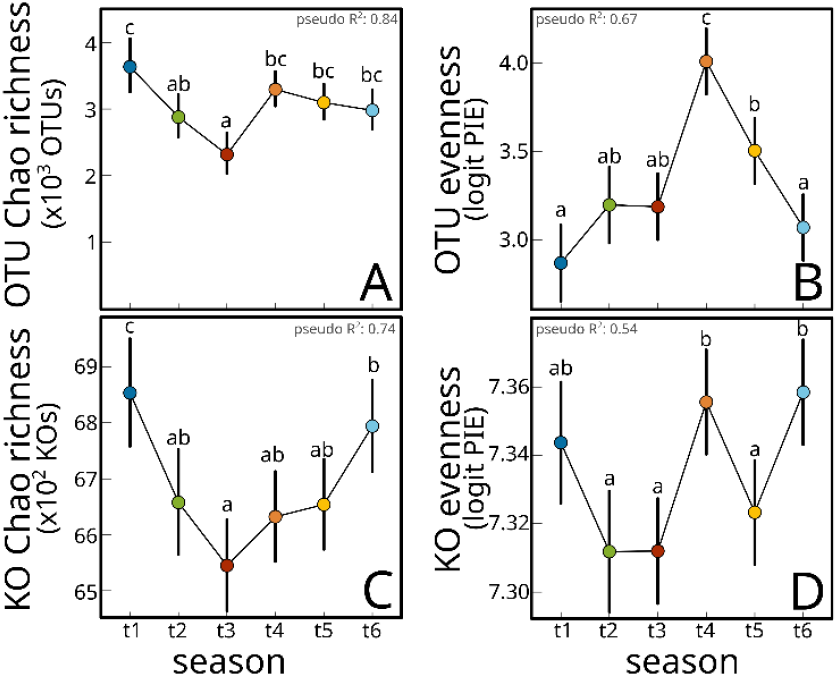
Estimated means of Chao OTU richness (**A**), OTU evenness (**B**), Chao KO richness (**C**) and KO evenness (**D**) from GLMs fitted on two diversity measures among the six season time points. Error bars show 95% confidence intervals. The Cox and Snell pseudo R^2^ is given in each corner of the respective plot. Significantly different time points within pair-wise comparisons in the post-hoc analysis are indicated by small letters.

### Core microbiota

Linear discriminant analysis identified several taxa as permanent biomarkers of *G. vermiculophylla* surfaces, i.e. they were at all times of the year significantly less characteristic for sediment and water samples (Fig. 5A, Table S6-7). The approach identified 8 OTUs that formed a permanent LEfSe-core of the algal host (Fig. 5A, Table S6), and also several higher taxonomic ranks as LEfSe-core groups (Table S7), including the phylum Bacteroidetes, the classes Alphaproteobacteria and Bacteroidea, four different orders (highest LDA score: Flavobacteriales), nine different families (highest LDA scores: Flavobacteriaceae and Hyphomonadaceae), and 12 different genera (Highest LDA score: *Granulosicoccus*). Altogether 69 OTUs formed a summer LEfSe-core of *G. vermiculophylla* and 33 OTUs formed a winter LEfSe core (Fig. 5B-C, Table S6). Biomarkers of summer (Fig. 5B, Table S7) were the dominant bacteria, the class Alphaproteobacteria, six orders (highest LDA scores: Rhodobacterales and Chitinophagales), eight families (highest LDA scores: Rhodobacteraceae and Saprospiraceae) and eight genera (highest LDA score: *Ulvibacter*). Taxonomic biomarkers of the winter season (Fig. 5C, Table S7) were the phyla Proteobacteria and Firmicutes, the class Clostridia, three orders (highest LDA score: Thiotrichales), six families (highest LDA score: Desulfobulbaceae) and 12 genera (highest LDA score: *Desulforhopalus*).

**Figure 5.**
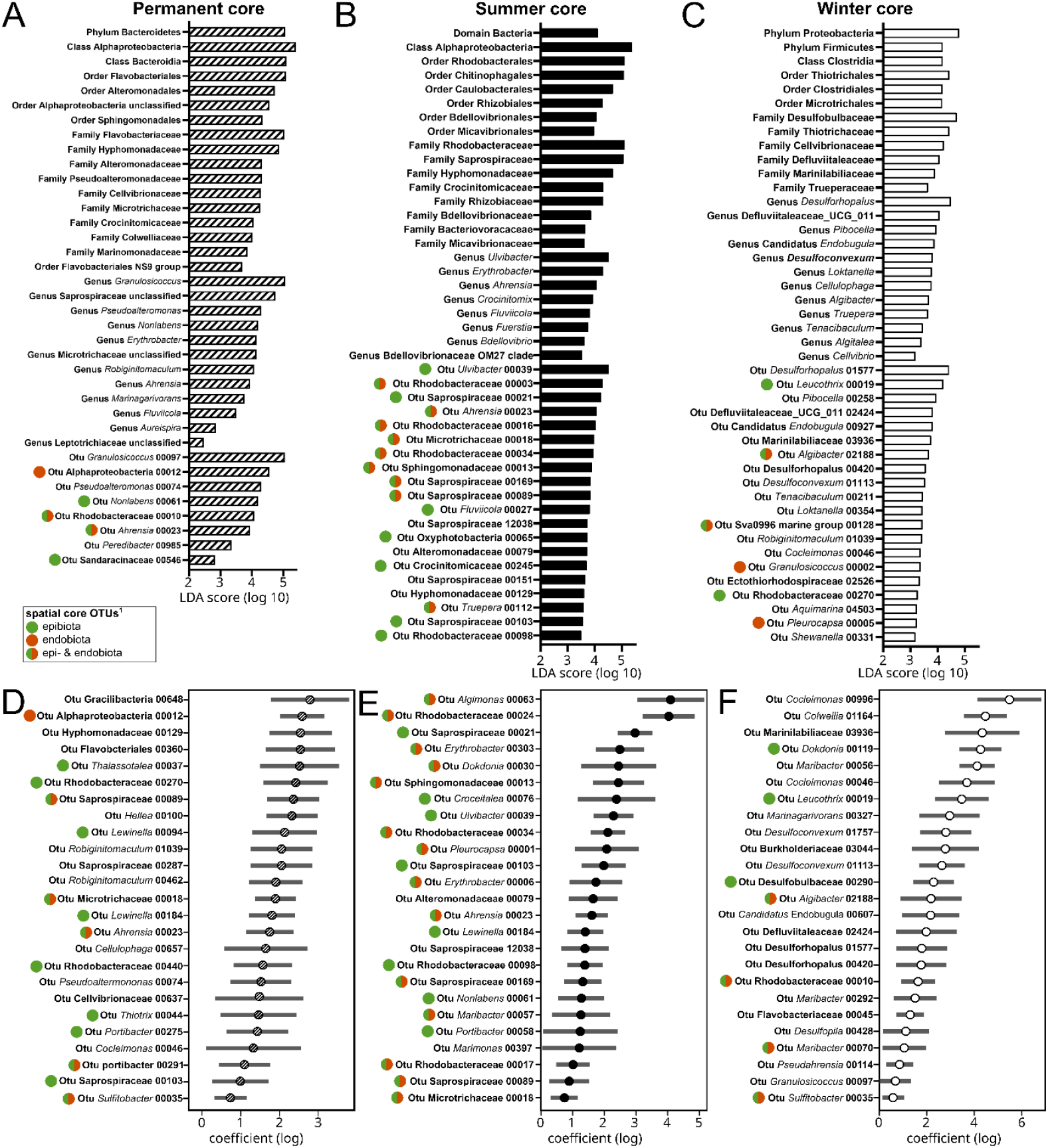
Permanent (**A, D**), summer (**B, E**) and winter (**C, F**) core epibiota associated with the rodophyte *Gracilaria vermiculophylla*. (**A-C**) Core taxa at different taxonomic levels detected with LEFse. (**D-F**) Core OTUs detected using mGLMs. Only the top 25 most abundant core OTUs are shown. ^(1)^ Green, red and green-red circles in front of the taxon labels indicate OTUs identified as spatial core OTUs in Bonthond *et al*. (2020).

The permanent and seasonal cores defined by the mGLM approach overlapped to some extent with the cores characterized by the linear discriminant analysis. The permanent mGLM-core counted 88 OTUs (Fig. 5D for the 25 most abundant OTUs, Table S6). 5 of the 8 OTUs LEfSe-core OTUs were also identified as part of the mGLM-core (Table S6). The summer mGLM-core of *G. vermiculophylla* counted 205 OTUs (Fig. 5E for the 25 most abundant OTUs, Table S6). 50 of the 69 the LEfSe summer core OTUs were also part of the summer mGLM-core. The mGLM winter core counted 285 OTUs. 21 out of the 33 LEfSe winter core OTUs were also part of the winter mGLM-core (Fig. 5F for the 25 most abundant OTUs, Table S6).

## DISCUSSION

This study revealed that epibiota associated with the seaweed holobiont *Gracilaria vermiculophylla* show strong seasonality. Prokaryotic composition and diversity are highly variable throughout the year. This variation was largely cyclic, showing similar trends over the three repetitive years in this study, with late winter and early summer as the extremes between which composition and diversity oscillated (Fig. 3-4). Our data also indicate that these seasonal shifts are not functionally redundant, as concurrent trends were found in terms of composition and diversity of predicted functions. Therewith, these findings support our hypothesis that the *G. vermiculophylla* holobiont has seasonal dynamics, providing evidence that seaweed associated epibiota undergo seasonal successional cycles. In line with this, we found numerous seasonal core taxa, that is, microbial OTUs or groups of higher taxonomic ranks, that were consistently associated with either summer or winter. In addition, our study also identified a permanent core, of microbial taxa which were consistently associated with the host independent of season.

### Temporal variation in epibiota is highly seasonal

Seasonal patterns have been described in seaweeds before, including chemical host processes such as metabolite production (Paix *et al*. 2019) or anti-fouling activity (Saha & Wahl 2013; Wang *et al*. 2018). Also in microbial communities associated with seaweeds, variability associated with seasonal changes has been observed (Bengtsson *et al*. 2010; Burgunter-Delamare *et al*. 2023; Korlević *et al*. 2021; Lachnit *et al*. 2011; Mancuso *et al*. 2016; Park *et al*. 2022; Tujula *et al*. 2010). However, while covering different seasons, the sampling in these studies is typically limited to one year (with the exception of Lachnit *et al*. 2011). Our findings are strongly in line with their observations, showing that also in the *G. vermiculophylla* holobiont, composition and diversity shift from one season to another. By repeating the seasonal sampling across three subsequent years, our study also resolves interannual trends which demonstrate that much of this temporal variation is repetitive and therefore truly seasonal (Fuhrman *et al*. 2015). Epibiota associated with *G. vermiculophylla*, are thus highly dynamic, but also resilient, as they undergo strong compositional shifts, but shift back towards compositions experienced in preceding years, in the same season.

Given the known structuring effects of both salinity (Stratil *et al*. 2014; Van Der Loos *et al*. 2023) and temperature on seaweed associated microbiota (Bonthond *et al*. 2023; Düsedau *et al*. 2023; Stratil *et al*. 2013), such seasonal environmental variables may well explain much of the here observed cyclic compositional and diversity changes. At the same time, they may also explain the pronounced differences in taxonomic composition that were observed between two sampling sites in the present study. However, besides the environment, also the host undergoes metabolic, physiological, and reproductive changes which can be season dependent (Liu *et al*. 2017 and references therein). Cycles in the host can also coincide with microbial life cycles, such as for example in the brown alga *Ascophyllum nodosum*, whose reproductive cycles are synchronized with the fungal symbiont *Stigmidium ascophylli* (Stanley 1992) or in *Acrochaetium* (Rhodophyta), in which bacterial metabolites (N-acyl-homoserine-lactones) regulate spore release (Weinberger *et al*. 2007). In this context, an interesting observation is that functional and taxonomic composition of the bacterial communities associated with *G. vermiculophylla* oscillated seasonally with similar intensity, whereas pronounced taxonomic differences between sites were hardly reflected by similar functional differences. Different *G. vermiculophylla* epibiota appear to be functionally similar between sites in a given season, but are functionally different among seasons. This suggests that the holobiont acquired season specific microbial functions, but is rather promiscuous to the microbes that provide them.

### Seasonal shifts in diversity

The shift from an OTU richness maximum in late winter to a minimum in summer (Fig. 4A), as well as an inverse pattern of evenness (Fig. 4B) was consistent across both populations and showed similarity to a study on temporal dynamics on the epibiota associated with the brown seaweed *Cystoseira compressa* (Mancuso *et al*. 2016). Moreover, this cyclic trend in Chao richness appeared to be even stronger in terms of predicted functions, which implies that seasonal changes in diversity are not functionally redundant, resulting in more diverse functions in winter in the associated epibiota. Hypothetically, a decrease in taxonomic and functional richness toward summer may be driven by rising temperatures and solar irradiance, with which metabolic rates increase (Clarke & Fraser 2004; Gillooly *et al*. 2001), and reinforce competition and extinction rates within the seaweed microbiota. If this is true, the seasonal diversity cycle in *G. vermiculophylla* may be a more general trend among seaweed holobionts. However, due to limited studies on seaweed holobionts including both seasonal and interannual samplings this remains to be evaluated in future studies, targeting different seaweeds.

### The core microbiome of Gracilaria vermiculophylla

Microbial cores have been studied across holobionts (Ainsworth *et al*. 2015; Burke *et al*. 2011; Schmitt *et al*. 2012; Shade & Handelsman 2012). Characterizing the core microbiota of a host, particularly over a large spatial or temporal scale (Shade & Handelsman 2012), provides an opportunity to detect patterns of stability and generality within a holobiont. After Bonthond *et al*. (2020) characterized a spatial core of epi- and endophytes within *G. vermiculophylla*, by sampling different populations of the host across its distribution range, the present work builds forward on this by characterizing temporal cores.

To identify core taxa of *G. vermiculophylla*, the present study utilized two compositional approaches (Shade & Handelsman 2012), resolving core OTUs based on statistically significant differential abundances. Both approaches corroborate our hypotheses that the *G. vermiculophylla* holobiont harbors prokaryotic taxa with strong temporal consistency, either associated permanently (Fig. 5A, D) or recurrently in summer (Fig. 5B, E) or winter (Fig. 5C, F).

Jointly, the spatial and temporal cores of Bonthond *et al*. (2020) and this study provide an elaborate, and to our knowledge unprecedented, impression of the core microbiota within a seaweed holobiont. A subset of 32 OTUs of the permanent mGLM-core was also identified as part of the spatial core in Bonthond *et al*. (2020). This set of OTUs is thus both geographically and temporally highly conserved, and represents a core that appears to be unconditionally present within this seaweed holobiont. In addition, 37 summer core OTUs and 50 winter core OTUs were also identified as spatial core OTUs in the previous study. While their presence in the *G. vermiculophylla* holobiont is season specific, they are also spatially and temporally consistent holobiont members.

Each of these taxa is of special interest, as their prevalent signal is unlikely coincidental. Perhaps most striking is the unclassified Alphaproteobacterial OTU (OTU12), which occurrence is 100% in both studies and whose poorly resolved identity may also indicate that the microbe is highly host-specific, and is difficult to isolate individually. Similarly, a member of the genus *Ahrensia* (OTU23), was present in 100% and > 99% of all samples in the spatial and present study, respectively. In addition to being identified as core spatial endophyte, both mGLM and LEfSe approaches resolved the *Ahrensia* OTU as permanent core member, implying a consistent presence of the holobiont.

The presence of a *Maribacter* core OTUs could hint at a host-microbe relationship within the *G. vermiculophylla* holobiont, similar to the chlorophyte *Ulva*, in which this bacterium plays a regulatory role in host morphogenesis (Weiss *et al*. 2017). Furthermore, the summer core also included the cyanobacterial OTU (OTU1), classified to *Pleurocapsa*, that was the most abundant OTU in the spatial study of Bonthond et al. (2020) and which was there found to be part of both epi- and endophytic cores. A closely related OTU, (OTU7, also classified to *Pleurocapsa*), also identified as spatial core endophyte and one of the most abundant OTUs in Bonthond *et al*. (2020), was here resolved as permanent core member. These cyanobacterial *Pleurocapsa* OTUs, are closely related to *Waterburya agarophytonicola*, which was isolated from the same host and has the genomic potential to synthesize various vitamins, including cobalamin (vitamin B_12_) for which *G. vermiculophylla* is auxotroph (Bonthond *et al*. 2021b). Such cyanobacterial core members may thus potentially play a role in vitamin acquisition for the seaweed host.

Noteworthy is the detection of three *Granulosicoccus* OTUs as part of the winter core (OTUs 2, 41 and 97), of which two were resolved as spatial core endophytes in Bonthond *et al*. (2020). The genus *Granulosicoccus* is considered to be a seaweed generalist and is often reported as core symbiont (Aires *et al*. 2023; Park *et al*. 2022). Metagenomic evidence from the Kelp *Nereocystis luetkeana* suggested that associated *Granulosicoccus* have diverse energy metabolism, but are incapable of autotrophic carbon fixation, which may indicate they obtain organic carbon from their seaweed host (Weigel *et al*. 2022). Moreover, Weigel *et al*. (2022) also found that *Granulosicoccus* have all genes necessary to synthesize cobalamin, which makes them another candidate vitamin source for the auxotrophic host *G. vermiculophylla*.

## CONCLUSION

Altogether, this study provides to the best of our knowledge one of the most detailed studies on seasonality in microbiota within a seaweed holobiont, with repeating a bi-monthly sampling over three years. Epibiota associated with *Gracilaria vermiculophylla* are dynamic, and seasonality drives much of this temporal variation in diversity and composition. These seasonal differences are likely linked to environmental conditions such as salinity and temperature, which fluctuate strongly throughout the year, especially in the shallow and intertidal habitats where this seaweed typically occurs. Despite strong compositional differences between North and Baltic Sea populations, similar cyclic patterns were resolved between the two populations, which reflects that despite strong differences between populations, they experience similar seasonal succession cycles, and which are thus likely natural to *G. vermiculophylla* holobionts. These succession cycles entail functional changes, as the cyclic trend was also evident in predicted functional composition. In contrast, differences between populations were minimal in terms of functional composition, which suggests that unlike the spatial shifts, seasonal changes are more functional. Based on this we posit that spatial variability in microbial composition within the *G. vermiculophylla* holobiont is more redundant than seasonal variability, because essential microbial functions can be obtained from a wide range of microbes.

While epibiota vary in space and time, we resolved 32 OTUs, which are permanent core members in this study and part of the spatial core characterized in earlier work of Bonthond *et al*. (2020). Therewith, this study demonstrates that certain microbial taxa are perpetual within the holobiont and are season and geography independent. This spatial and temporal core presents a subset of candidate microorganisms that may play important roles in host functioning and which merits future attention.

## AUTHOR CONTRIBUTIONS

CMM, FW and GB conceptualized the study. Field collections were conducted by CMM, FW, LD and GB. CMM, FW, LD, MG, SK and GB conducted laboratory work. CMM, FW and GB processed data. CMM, FW and GB conducted the formal analysis. CMM, FW and GB drafted the manuscript. All authors contributed to writing and revising the manuscript.

## ACKNOWLEDGEMENTS

This study was funded by the Deutsche Forschungsgemeinschaft (DFG), through a grant awarded to FW (DFG grant number WE2700/5-1). The authors are grateful to Nabila Elarbi, Barbora Burýšková and Fatiha Kalam Nisa for support during fieldwork and to the Deutsche Bundesamt für Seeschifffahrt und Hydrographie (BSH) for kindly providing temperature data.

## DATA AVAILABILITY STATEMENT

The raw de-multiplexed V4-16S rDNA gene amplicon reads and associated metadata are available from the SRA database under the Bioproject accession number PRJNA1155875. Other data and R-scripts for analyses are available on GitHub at https://github.com/gbonthond/Seasonalilty_seaweed_holobiont.

## SUPPORTING INFORMATION

**Table S1**. PERMANOVA community composition all substrates

**Table S2**. PERMANOVA community composition on only algal substrate

**Table S3**. PERMANOVA predicted functional composition on only algal substrate

**Table S4. (A)** ANOVA table for asymptotic richness (S_Chao_) based on OTUs and KOs. **(B)** Post-hoc pair-wise comparisons within the factor season (t1 – t6) for asymptotic richness (S_Chao_) based on OTUs and KOs

**Table S5. (A)** ANOVA table for evenness (Probability of Interspecific Encounter) based on OTUs and KOs. **(B)** Post-hoc pair-wise comparisons within the factor season (t1 – t6) for for evenness (Probability of Interspecific Encounter) based on OTUs and KOs.

**Table S6**. Epiphytic cores

**Table S7**. Higher taxonomic rank cores

**Figure S1**. Stacked bar plots showing community composition by substrate

**Figure S2**. Stacked bar plots showing community composition by population

**Figure S3**. Stacked bar plots showing community composition by population, year, and collection event

**Figure S4**. nMDS with algal, water, and sediment samples

