## Supporting Information for "Seasonal cycles in a seaweed holobiont: A multiyear time series reveals repetitive microbial shifts and core taxa"

### FIGURES

**Figure S1.** Stacked bar plots showing community composition by substrate
**Figure S2.** Stacked bar plots showing community composition by population
**Figure S3.** Stacked bar plots showing community composition by population, year and collection event
**Figure S4.** nMDS with sediment and water samples

### TABLES

**Table S1.** PERMANOVA community composition all substrates
**Table S2.** PERMANOVA community composition on only algal
**Table S3.** PERMANOVA predicted functional composition
**Table S4.** Analysis of Deviance tables from diversity metrics analyzed with GLMs and likelihood ratio tests.
**Table S5.** Table S5. Post-hoc pairwise comparisons for all levels within the predictor season.
**Table S6.** Epiphytic cores
**Table S7.** Higher taxonomic rank cores

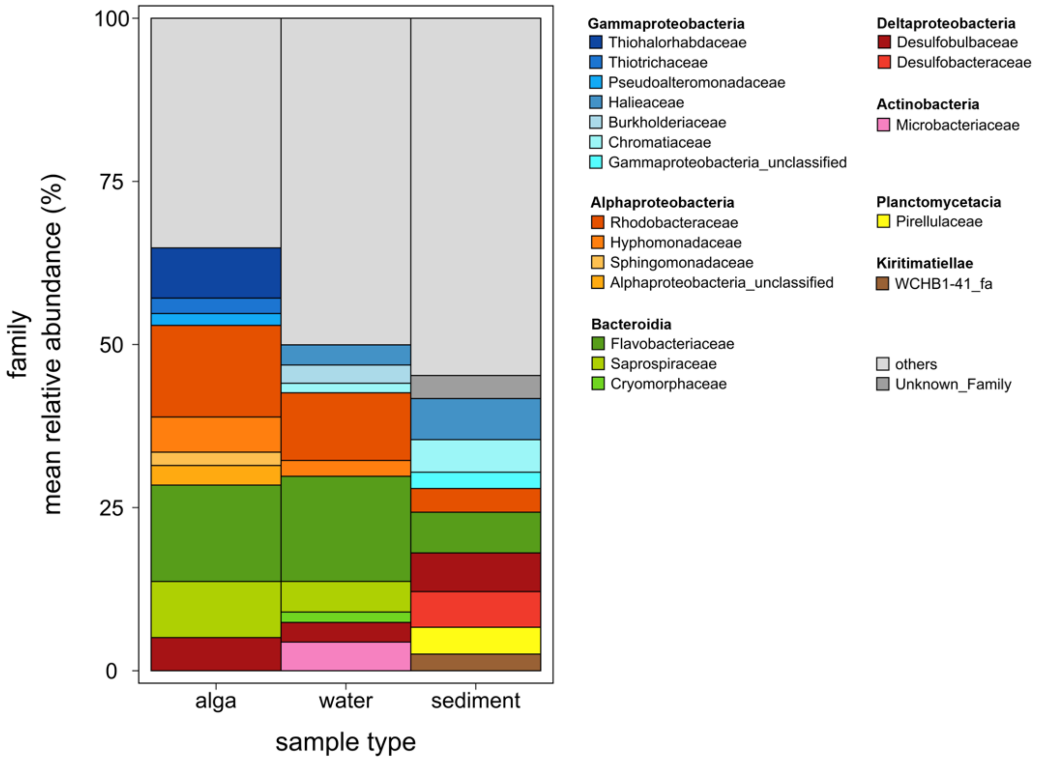


**Figure S1.** Microbial community composition of the ten most abundant families associated with the sample types alga, water and sediment. Shown is the mean relative abundance in percent (%). Main colours fit to the taxonomic class and corresponding families are illustrated with a variation of that main colour. The fraction “others” counts all families not represented in the ten most abundant members. Normalized data on total OTU read counts is shown.

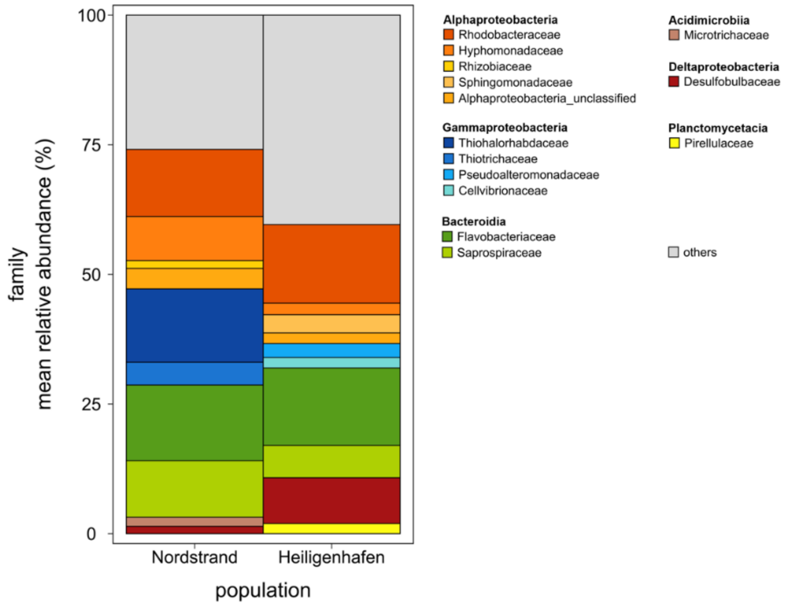


**Figure S2.** Microbial community composition of the ten most abundant families associated with the surface of the red seaweed *Gracilaria vermiculophylla* divided between the two populations Nordstrand (North Sea) and Heiligenhafen (Baltic Sea). Shown is the mean relative abundance in percent (%). Main colours fit to the taxonomic class and corresponding families are illustrated with a variation of that main colour. The fraction “others” counts all families not represented in the ten most abundant members. Normalized data on total OTU read counts is shown.

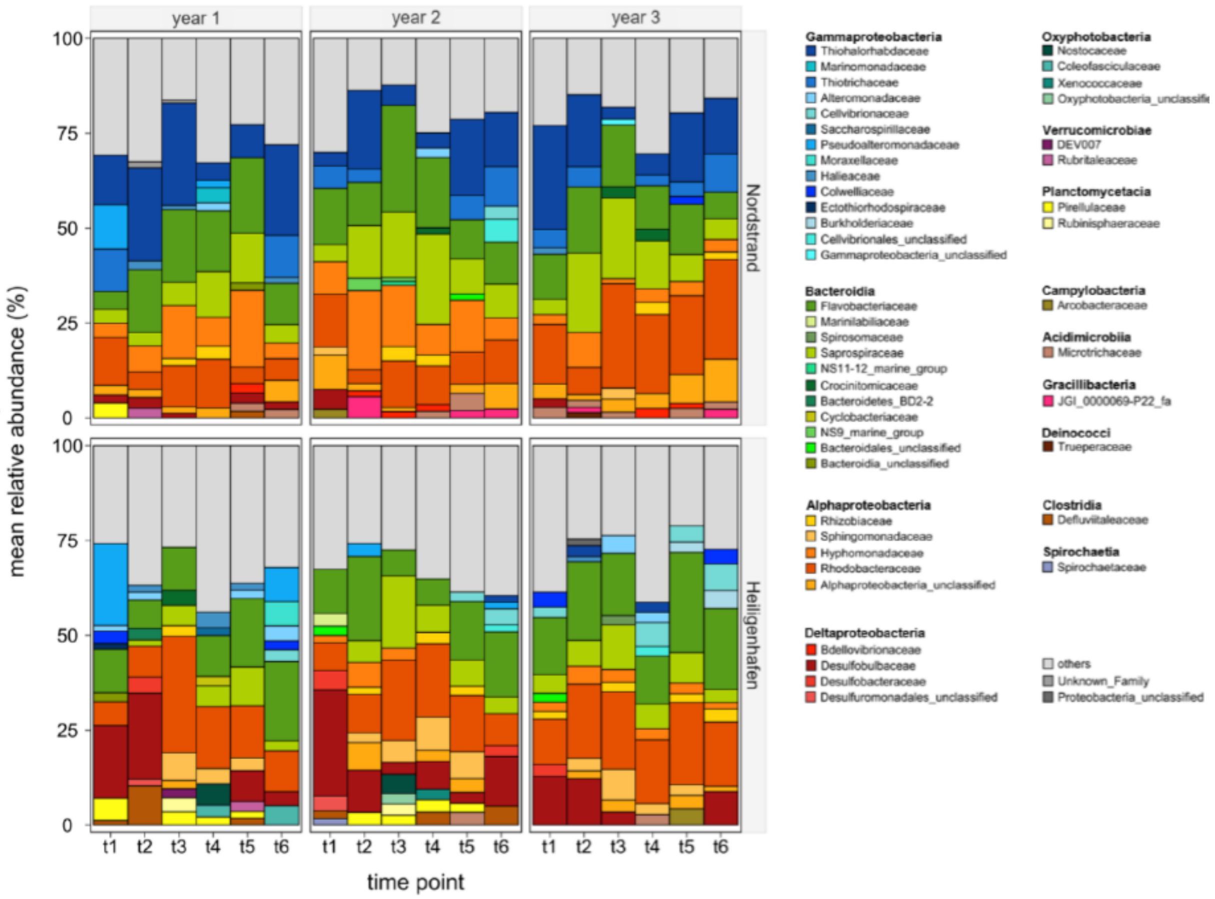


**Figure S3.** Microbial community composition of the most abundant families associated with the surface of the red seaweed *Gracilaria vermiculophylla*, by population, year and collection event. Sizes of the stacked bars are relative percentages (%), averaged over the replicates within the same group.

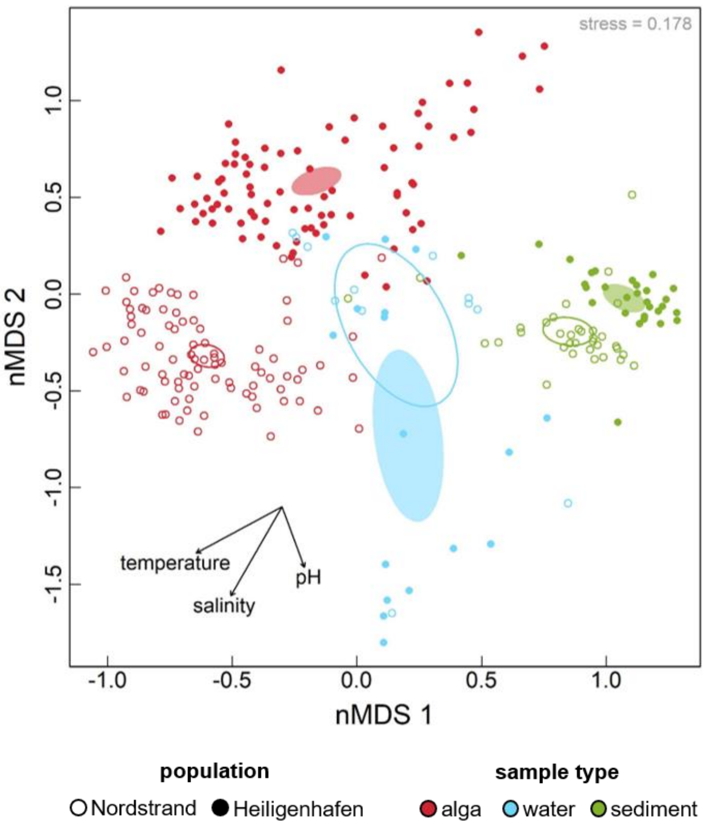


**Figure S4**. **nMDS with algal, water, and sediment samples.** Microbial community composition associated with the red seaweed *Gracilaria vermiculophylla* among three different sample types. The sample types alga (red), water (blue), and sediment (green) are pooled over the three collection years. The Nordstrand (North Sea) population is visualized with empty and Heiligenhafen (Baltic Sea) with filled dots and ellipses, respectively. The ellipses are represented with a 95% confidence interval. Additionally, the abiotic factors temperature, salinity, and pH are plotted. The stress value is given in the upper right corner. The microbial diversity is represented by the rarefied OTU read counts.

**Table S1.** PERMANOVA table for the microbial composition within all substrates (alga, water, sediment).

| | *df* | *Sums Sqs.* | *Mean Sqs.* | *F-value* | $R^2$ | *p-value*[1] |
|---|---|---|---|---|---|---|
| LSD | 1 | 10.918 | 10.918 | 57.982 | 0.107 | **< 0.001** |
| season | 5 | 4.689 | 0.938 | 4.980 | 0.046 | **< 0.001** |
| year | 2 | 1.548 | 0.774 | 4.111 | 0.015 | **< 0.001** |
| population | 1 | 5.416 | 5.416 | 28.762 | 0.053 | **< 0.001** |
| substrate | 2 | 11.909 | 5.955 | 31.622 | 0.117 | **< 0.001** |
| season:year | 10 | 4.507 | 0.451 | 2.394 | 0.044 | **< 0.001** |
| season:population | 5 | 3.410 | 0.682 | 3.621 | 0.033 | **< 0.001** |
| year:population | 2 | 1.250 | 0.625 | 3.320 | 0.012 | **< 0.001** |
| season:substrate | 9 | 4.717 | 0.524 | 2.783 | 0.046 | **< 0.001** |
| year:substrate | 4 | 2.280 | 0.570 | 3.027 | 0.022 | **< 0.001** |
| population:substrate | 2 | 3.893 | 1.947 | 10.337 | 0.038 | **< 0.001** |
| season:year:population | 10 | 3.839 | 0.384 | 2.039 | 0.038 | **< 0.001** |
| season:year:substrate | 14 | 4.473 | 0.320 | 1.697 | 0.044 | **< 0.001** |
| season:population:substrate | 8 | 3.001 | 0.375 | 1.992 | 0.029 | **< 0.001** |
| year:population:substrate | 4 | 1.545 | 0.386 | 2.051 | 0.015 | **< 0.001** |
| season:year:population:substrate | 7 | 1.788 | 0.255 | 1.357 | 0.018 | 0.002 |
| Residuals | 175 | 32.953 | 0.188 | | 0.323 | |
| Total | 261 | 102.136 | | | 1 | |

*[1]Terms were sequentially tested (from first to last) and p-values were obtained with 9999 permutations*
*Abbreviations: Permutational Analysis of Variance (PERMANOVA), Log-transformed sequencing depth (LSD)*

**Table S2.** PERMANOVA table for the microbial composition solely within the substrate alga is shown.

| | *df* | *Sums Sqs.* | *Mean Sqs.* | *F-value* | $R^2$ | *p-value*[1] |
|---|---|---|---|---|---|---|
| LSD | 1 | 7.206 | 7.206 | 37.777 | 0.118 | **< 0.001** |
| season | 5 | 5.596 | 1.119 | 5.867 | 0.092 | **< 0.001** |
| year | 2 | 1.676 | 0.838 | 4.393 | 0.028 | **< 0.001** |
| population | 1 | 6.659 | 6.659 | 34.907 | 0.109 | **< 0.001** |
| season:year | 10 | 4.653 | 0.465 | 2.439 | 0.076 | **< 0.001** |
| season:population | 5 | 3.964 | 0.793 | 4.156 | 0.065 | **< 0.001** |
| year:population | 2 | 1.515 | 0.758 | 3.972 | 0.025 | **< 0.001** |
| season:year:population | 10 | 4.260 | 0.426 | 2.233 | 0.070 | **< 0.001** |
| Residuals | 133 | 25.371 | 0.191 | | 0.417 | |
| Total | 169 | 60.900 | | | 1 | |

*[1]Terms were sequentially tested (from first to last) and p-values were obtained with 9999 permutations*
*Abbreviations: Permutational Analysis of Variance (PERMANOVA), Log-transformed sequencing depth (LSD)*

**Table S3.** PERMANOVA table for the predicted functional composition solely within the substrate alga is shown.

| | *df* | *Sums Sqs.* | *Mean Sqs.* | *F-value* | $R^2$ | *p-value*[1] |
|---|---|---|---|---|---|---|
| LSD | 1 | 17.036 | 17.036 | 237.475 | 0.544 | **< 0.001** |
| season | 5 | 1.136 | 0.227 | 3.167 | 0.036 | **0.001** |
| year | 2 | 0.499 | 0.250 | 3.478 | 0.016 | **0.008** |
| population | 1 | 0.214 | 0.214 | 2.979 | 0.007 | **0.041** |
| season:year | 10 | 1.142 | 0.114 | 1.591 | 0.036 | **0.043** |
| season:population | 5 | 0.438 | 0.088 | 1.220 | 0.014 | 0.254 |
| year:population | 2 | 0.110 | 0.055 | 0.770 | 0.004 | 0.553 |
| season:year:population | 10 | 1.223 | 0.122 | 1.705 | 0.039 | **0.024** |
| Residuals | 133 | 9.541 | 0.072 | | 0.304 | |
| Total | 169 | 31.338 | | | 1 | |

*[1]Terms were sequentially tested (from first to last) and p-values were obtained with 9999 permutations*
*Abbreviations: Permutational Analysis of Variance (PERMANOVA), Log-transformed sequencing depth (LSD)*

**Table S4A.** ANOVA table for asymptotic richness ($S_{Chao}$) based on OTUs and KOs.

| | *df* | *OTUs* | | *KOs* | |
|---|---|---|---|---|---|
| | | *LR Chisq.* | *p-value* | *LR Chisq.* | *p-value* |
| LSD | 1 | 269.792 | **< 0.001** | 192.504 | **< 0.001** |
| season | 5 | 59.076 | **< 0.001** | 34.026 | **< 0.001** |
| year | 2 | 97.452 | **< 0.001** | 64.126 | **< 0.001** |
| population | 1 | 15.163 | **< 0.001** | 1.324 | 0.2498 |
| season:year | 10 | 53.052 | **< 0.001** | 21.214 | **0.01965** |
| season:population | 5 | 40.330 | **< 0.001** | 9.536 | 0.08949 |
| year:population | 2 | 8.776 | 0.012 | 3.872 | 0.14430 |
| season:year:population | 10 | 19.564 | 0.034 | 20.689 | **0.02337** |

*Abbreviations: Log-transformed sequencing depth (LSD)*

**Supplementary Table S4B.** Post-hoc pair-wise comparisons within the factor season (t1 – t6) for asymptotic richness ($S_{Chao}$) based on OTUs and KOs.

| *Contrast* | *df* | *OTUs* | | *KOs* | |
|---|---|---|---|---|---|
| | | *t-value* | *p-value* | *t-value* | *p-value* |
| t1:t2 | 133 | 2.937 | **0.044** | 2.721 | 0.0393 |
| t1:t3 | 133 | 5.142 | **< 0.001** | 5.003 | **< 0.001** |
| t1:t4 | 133 | 1.393 | 0.731 | 3.379 | **0.0066** |
| t1:t5 | 133 | 2.244 | 0.225 | 3.057 | **0.0166** |
| t1:t6 | 133 | 2.668 | 0.089 | 0.157 | 0.9238 |
| t2:t3 | 133 | 2.454 | 0.146 | 1.991 | 0.4575 |
| t2:t4 | 133 | -1.921 | 0.394 | 0.476 | 0.9985 |
| t2:t5 | 133 | -1.018 | 0.911 | 0.107 | 1.0000 |
| t2:t6 | 133 | -0.451 | 0.998 | -2.668 | 0.2460 |
| t3:t4 | 133 | -4.425 | **< 0.001** | -1.636 | 0.6575 |
| t3:t5 | 133 | -3.585 | **0.006** | -2.048 | 0.4001 |
| t3:t6 | 133 | -2.974 | **0.040** | -5.037 | **0.0004** |
| t4:t5 | 133 | 1.026 | 0.908 | -0.401 | 0.9989 |
| t4:t6 | 133 | 1.537 | 0.641 | -3.382 | 0.0615 |
| t5:t6 | 133 | 0.580 | 0.992 | -3.019 | 0.1437 |

**Table S5A.** ANOVA table for evenness (Probability of Interspecific Encounter) based on OTUs and KOs.

| | *Df* | ***OTUs*** | | ***KOs*** | |
|---|---|---|---|---|---|
| | | ***LR Chisq.*** | ***p-value*** | ***LR Chisq.*** | ***p-value*** |
| season | 5 | 86.574 | **< 0.001** | 39.519 | **< 0.001** |
| year | 2 | 8.234 | **0.01629** | 7.628 | **0.02206** |
| population | 1 | 63.915 | **< 0.001** | 19.463 | **< 0.001** |
| season:year | 10 | 36.697 | **< 0.001** | 22.587 | **0.01238** |
| season:population | 5 | 30.198 | **< 0.001** | 25.808 | **< 0.001** |
| year:population | 2 | 0.964 | 0.61755 | 2.265 | 0.32221 |
| season:year:population | 10 | 40.915 | **< 0.001** | 40.085 | **< 0.001** |

**Supplementary Table S5B.** Post-hoc pair-wise comparisons within the factor season (t1 – t6) for for evenness (Probability of Interspecific Encounter) based on OTUs and KOs.

| ***Contrast*** | ***df*** | ***OTUs*** | | ***KOs*** | |
|---|---|---|---|---|---|
| | | ***t-value*** | ***p-value*** | ***t-value*** | ***p-value*** |
| t1:t2 | 133 | -2.214 | 0.2385 | 2.865 | 0.0536 |
| t1:t3 | 133 | -2.327 | 0.1906 | 3.062 | **0.0312** |
| t1:t4 | 133 | -8.109 | **< 0.001** | -0.764 | 0.9729 |
| t1:t5 | 133 | -4.622 | **< 0.001** | 2.073 | 0.3078 |
| t1:t6 | 133 | -1.466 | 0.6864 | -1.126 | 0.8697 |
| t2:t3 | 133 | 0.073 | 1.0000 | -0.041 | 1.0000 |
| t2:t4 | 133 | -5.672 | **< 0.001** | -3.659 | **0.0048** |
| t2:t5 | 133 | -2.169 | 0.2593 | -0.982 | 0.9229 |
| t2:t6 | 133 | 0.903 | 0.9452 | -4.028 | **0.0013** |
| t3:t4 | 133 | -6.235 | **< 0.001** | -3.936 | **0.0018** |
| t3:t5 | 133 | -2.442 | 0.1495 | -1.026 | 0.9086 |
| t3:t6 | 133 | 0.904 | 0.9449 | -4.345 | **< 0.001** |
| t4:t5 | 133 | 3.836 | **0.0026** | 2.921 | **0.0461** |
| t4:t6 | 133 | 7.131 | **< 0.001** | -0.364 | 0.9991 |
| t5:t6 | 133 | 3.347 | 0.0133 | -3.319 | 0.0145 |

**Table S6.** See separate excel file

**Table S7.** See separate excel file
